# Early Detection of Neuroinflammation and White Matter Damage Following Dorsal Spinal Nerve Root Sectioning in a Nonhuman Primate Model

**DOI:** 10.1101/2025.07.02.662878

**Authors:** Feng Wang, John C. Gore, Li Min Chen

## Abstract

**Purpose:** Dorsal rhizotomy, or spinal dorsal nerve root lesioning, is a surgical procedure used to treat intractable nerve pain by selectively severing sensory afferent nerve roots. This study aimed to evaluate whether multiparametric MRI, including diffusion tensor imaging (DTI), quantitative magnetization transfer (qMT), and chemical exchange saturation transfer (CEST), can sensitively detect structural and biochemical changes in the intact spinal cord following a focal dorsal nerve root section in a non-human primate model.

**Methods:** In four squirrel monkeys, unilateral dorsal nerve roots at cervical segments C4 and C5 were surgically transected. MRI data were collected using a 9.4 T scanner with a custom saddle-shaped transmit-receive quadrature coil before and one week after lesioning. DTI-derived fractional anisotropy (FA), axial diffusivity (AD), and radial diffusivity (RD); qMT-derived pool size ratio (PSR); and CEST and nuclear Overhauser enhancement (NOE) effects were quantified across seven spinal cord regions of interest (ROIs). CEST and NOE effects were extracted using five-pool Lorentzian fitting of Z-spectra.

**Results:** At the lesioned dorsal nerve root bundles, FA, PSR, and NOE (-1.6 ppm) values decreased, while RD and CEST (3.5 ppm) increased, consistent with fiber degeneration, demyelination, and inflammation. Similar, though less pronounced, changes were observed in the dorsal root entry zone, particularly within the first week post-lesion.

**Conclusion:** Multiparametric MRI enables region-specific detection of early spinal cord pathology as soon as one week following dorsal nerve root injury. These findings support its potential as a noninvasive tool for monitoring secondary degeneration due to spinal nerve damage and for evaluating outcomes of therapeutic interventions.

## 1 INTRODUCTION

Traumatic nerve root injuries typically occur in individuals who have experienced major trauma, such as high-velocity accidents. Cervical injuries are most common due to the spine’s mobility. These injuries can also arise from spinal dorsal nerve root lesioning (dorsal rhizotomy), neurosurgical procedure in which the sensory afferent nerve rootlets of the dorsal spinal roots are selectively severed to reduce pathological sensory inputs.^1^ This procedure is used in children with cerebral palsy, individuals suffering from intractable cancer or neuropathic pain, and patients with spinal cord injury (SCI)-related pain or spasticity.^2^ Given the anatomical and physiological similarity between non-human primates (NHPs) and humans, NHPs serve as realistic and translationally relevant preclinical models for studying nerve root injuries and the resulting secondary spinal cord pathology. In a previous NHP model of traumatic SCI, we demonstrated that while functional disruption can be assessed using functional MRI, structural and biochemical changes within the injured spinal cord are better evaluated using multiparametric MRI techniques. These approaches offer detailed insights into underlying pathological processes. Unlike SCI, dorsal nerve root sectioning occurs outside the spinal cord. However, the disruption of nerve bundles frequently leads to secondary pathological changes within the spinal cord, which may contribute to or mediate the relief of clinical symptoms. In this study, we aimed to characterize those structural and biochemical changes using multiparametric MRI in an NHP model involving targeted unilateral dorsal nerve root transection at two cervical spinal cord segments (C4 and C5). We hypothesized that focal, selective unilateral dorsal nerve root sectioning would initiate afferent fiber degeneration, potentially leading to secondary injury and tissue damage extending into the central branches of the affected nerve root bundles and their entering zones – specifically, the dorsal root entry zone (DREZ).

Quantitative assessment of SCI remains challenging. Clinical diagnosis of spinal cord and peripheral nerve injuries primarily relies on anatomical MRI with varying contrasts.^3, 4^ This limitation is partly due to technical constraints such as the small size of the cord and nerves, local magnetic field inhomogeneity, relatively low signal-to-noise ratio (SNR), restricted scan time, and motion artifacts from cerebrospinal fluid (CSF) pulsations associated with cardiac and respiratory cycles.^5, 6^ Currently, diffusion tensor imaging (DTI) is the most widely used quantitative MRI technique due to its relatively high reliability and straightforward data acquisition. DTI characterizes the anisotropy of water diffusion in white matter (WM) using a three-dimensional tensor model, generating metrics that reflect microstructural features such as fiber density, orientation, and integrity.^7, 8^ Advances in MRI hardware and imaging sequences, have enabled DTI to be applied to various spinal pathologies, including traumatic injury.^9, 10^ In addition, quantitative MT (qMT) imaging is capable of detecting axonal demyelination and fiber loss, while also identifying abnormal tissue changes adjacent to injury sites. Chemical exchange saturation transfer (CEST) imaging provides sensitivity to molecular composition, offering additional information about lesion-related metabolic and biochemical alterations, particularly in the early stages of injury. In recent years, we have leveraged the capabilities of high-field MRI to integrate DTI, qMT, and CEST in a single session for evaluating tissue composition and biochemical changes associated with SCI in NHPs subjected to controlled dorsal column transection.^11–18^ We have also extended this approach to rodent models of lumbar contusion injury.^19–21^ This multiparametric strategy has enabled comprehensive assessment of SCI progression and recovery, and has supported the identification of mechanism-informed MRI biomarkers.

This study applied our established multiparametric MRI approach to examine regional changes in the injured dorsal nerve root bundles and the adjacent intact spinal cord following dorsal root nerve transection in individual squirrel monkeys. The central aim was to investigate how damage to nerve root fibers located outside the spinal cord influences spinal cord integrity, particularly at the dorsal root–spinal cord interface, where no direct trauma occurs. We also evaluated whether DTI, qMT, and CEST can detect subtle pathological changes at the early stage (one week) after nerve root sectioning. The insights gained from this study will help advance the use of multiparametric MRI in detecting and characterizing early and localized spinal cord pathology resulting from clinically relevant nerve root injuries.

## 2 METHODS

### 2.1 Dorsal nerve root transection

Four adult male squirrel monkeys (*Saimiri sciureus,* 6-8 years old) were studied. The cervical segments C4/5-C6/7 receive sensory input from the upper arm and hand. This injury model is designed to selectively and unilaterally disrupt dorsal nerve roots of C4 and C5 segments. Dorsal nerve roots at the C4 and C5 segments were sectioned for squirrel monkeys. In brief, under surgical level anesthesia and aseptic conditions, the dorsal portion of the cervical spinal cord was exposed. 7 brunches of dorsal nerve roots were transected on one side before they enter the spinal cord using fine surgical scissors at both C4 and C5 levels. Each of the 7 nerve roots was separately cut and fully disconnected. The dura was replaced with a small piece of gelfilm, and the wound was enclosed. All procedures were approved by the Institutional Animal Care and Use Committee (IACUC) at Vanderbilt University and adhered to the NIH guidelines for the care and use of laboratory animals.

### 2.2 *In vivo* MRI data acquisition

MRI data were acquired from the cervical spinal cords before and after injury. MRI data included in this work were acquired around week 1 after injury to identify features at the early stage. During MRI data acquisitions, each monkey was anesthetized with isoflurane (0.8-1.2%), delivered in a 70:30 N_2_O/O_2_ mixture, and mechanically ventilated. Vital physiological signs were continuously monitored and maintained at stable levels throughout the imaging session.

All MR images were acquired using a customized saddle-shaped transmit-receive quadrature surface coil (each of the two loops is 30×30 mm^2^, mounted on 45-mm-diameter cylindrical surface) positioned around the cervical spine region at 9.4T. This coil was optimized to sample data from cervical spinal cord.^22^ It reduced unwanted signals from distant moving tissues and increases SNR. Magnetization transfer contrast (MTC)-weighted structural images were acquired in multiple orientations with enhanced contrast,^11^ facilitating the precise slice placement of DTI, qMT, and CEST (Fig. 1B).

**Figure 1.**
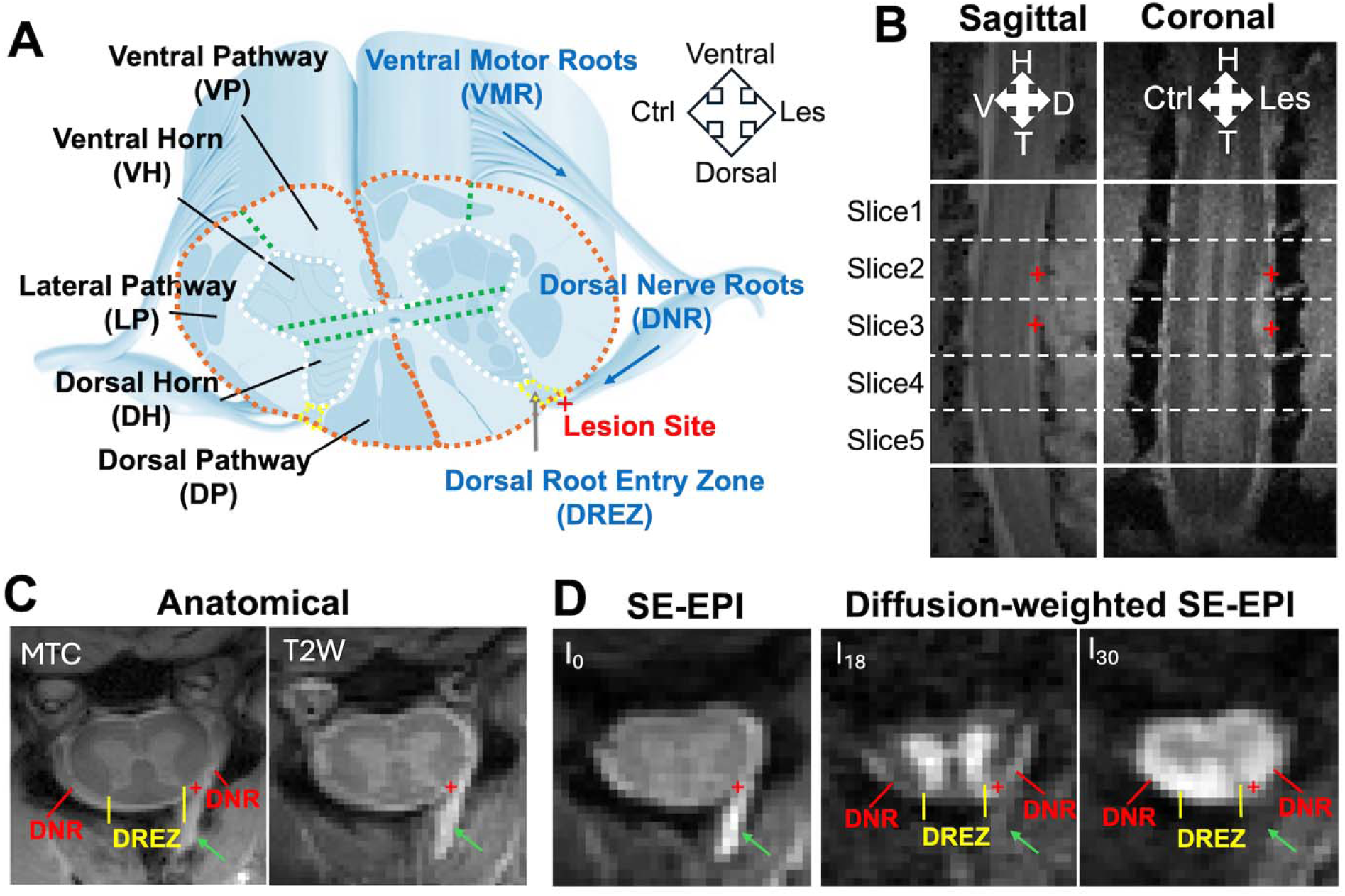
Anatomical images after unilateral dorsal nerve root lesion. (A) Schematic plot showing the regions of spinal cord and lesion site (indicated by ‘+’ at dorsal nerve roots). White matter regions (VP, ventral pathway; LP, lateral pathway; DP, dorsal pathway) and gray matter regions (VH, ventral horn; DH, dorsal horn) are separated by dashed lines. The dorsal root entry zones (DREZ) were highlighted by yellow dashed line. Ctrl, contralateral; Les, lesion. (B) MRI images with magnetization transfer contrast (MTC) in sagittal and coronal orientations. ‘+’ symbols indicate the targeted lesion sites on dorsal sensory nerve roots C4 and C5. H, head; T, tail; Ctrl, contralateral; Les, lesion. (C) High-resolution anatomical MTC and T_2_-weighted (T2W) images. (D) Spin-echo echo-planar image (SE-EPI). I_0_ is non-diffusion-weighted SE-EPI image, I_18_ and I_30_ are selected diffusion-weighted SE-EPI images, with diffusion gradients more parallel and more perpendicular to the cord respectively. The green arrows indicate fluids accumulated after lesion. The red lines indicate dorsal nerve root (DNR) bundles. The yellow lines indicate DREZ. The images were from one representative subject after lesion.

The DTI data acquisition was centered around the lesion level C4-C5 (Fig. 1). The diffusion-weighted spin-echo sequence used an echo planar imaging readout (TR/TE = 3000/33 ms, 4 shots, resolution = 0.333×0.333 mm^2^, slice thickness = 3 mm, 5 slices), with *b* values of 1000 s/mm^2^, sampling 30 directions (equally spaced). The qMT data were collected using a 2D MT-weighted spoiled gradient echo sequence (TR 24 ms, flip angle =7°, matrix size 128×128, 4 acquisitions, resoltuion = 0.25×0.25×3 mm^3^) for each spinal cord segment. Gaussian-shaped saturation pulses (θ_sat_ = 220° and 820°, pulse width = 10 ms, 10 RF offsets with a constant logrithmic interval between 1 and 100 kHz) were used. CEST and nuclear Overhauser enhancement (NOE) measurements were performed using a 5s continuous wave (CW) saturation of amplitude 1.0 μT followed by a spin-echo echo-planar-imaging acquisition (TR = 7.5 s, TE = 17.6 ms, 2 shots, matrix of 64×64, resolution = 0.5×0.5×3 mm^3^). Z-spectra were acquired with RF offsets from -2000 to 2000 Hz (-5 to 5 ppm at 9.4T) with an interval of 80 Hz (0.2 ppm at 9.4T). Reference scans were obtained at the beginning and the end of each acquisition using an RF offset at 100 kHz.

High-resolution MTC and T_2_-weighted (T2W) anatomical images (Fig. 1C) were acquired at 0.125 x 0.125 mm^2^ and 0.250 x 0.250 mm^2^ respectively, using the same geometry as the quantitative MRI. The total acquisition time for the above MRI data was approximately 1 hour.

### 2.3 MRI Data Analyses

MRI data were analyzed using MATLAB (The MathWorks). Conventional DTI parameters, including fractional anisotropy (FA), axial diffusivity (AD), and radial diffusivity (RD), were quantified using the DTI Toolbox^23^ based on diffusion data from the *b* = 1000 s/mm^2^ shell. The non-diffusion-weighted SE-EPI (I_0_) images were diffeomorphically registered slice-wise to the high-resolution MTC images, and all other diffusion weighted volumes (I_1_-I_30_) were registered to I_0_ image using a rigid registration algorithm (Fig. 1).

The qMT parameters including pool size ratio (PSR) were calculated using the 2-pool (free water and bound water pools) Ramani model.^12^ CEST images were acquired using a 5s continuous wave (CW) saturation of amplitude 1.0 µT (51 RF offsets equally spaced from -5 ppm to 5 ppm) followed by a spin-echo echo-planar-imaging acquisition (TR/TE = 7500/18 ms, 2 shots). The amplitudes of CEST and NOE effects at RF offsets -3.5, -1.6, 0, 2.0 and 3.5 ppm from multiple proton pools were quantified using a five-pool Lorentzian fitting of each Z-spectrum.^14^

High-resolution MTC, T2W and diffusion-weighted images were used as references for the manual selection of regions of interest (ROIs) for quantification, based on the known anatomy and landmarks in the monkey.^24^ Seven ROIs were manually delineated along the regional boundaries using MATLAB, including WM in lateral pathway (LP), ventral pathway (VP), dorsal pathway (DP), dorsal root entry zone (DREZ), and dorsal nerve roots (DNR), and gray matter (GM) in ventral horn (VH) and dorsal horn (DH).

### 2.4 Statistical Analysis

A two-sided Wilcoxon rank sum test was performed to obtain *p* value and evaluate the statistical significance of differences in affected C4 and C5 segments across subjects (number of injured segments = 8) for each individual ROI, comparing the lesion and contralateral non-lesioned side with pre-lesion healthy tissues. A *p*-value < 0.05 was considered statistically significant.

## 3 RESULTS

### 3.1 Nerve root injury is visible on anatomical MTC and diffusion-weighted images

One week after unilateral dorsal root nerve sectioning, MTC and T2W images clearly revealed the injury levels and affected spinal cord segments (Fig. 1). The seven nerve bundles at two cervical levels were transected before entering the spinal cord at near-horizontal angles (Fig. 1A). As a results, hyperintensities appeared along the dorsal surface of the spinal cord in both sagittal and coronal MTC images (Fig. 1B). In axial MTC and T2W images, the accumulated fluids appeared hyperintense (green arrows in Fig. 1C). The dorsal nerve root (DNR) bundles were well visualized in the high-resolution MTC image (Fig. 1C). In diffusion-weighted images, fluid signals were much darker than in non-diffusion-weighted images (green arrows in Fig. 1D), which enhanced the visibility of DNR bundles. The DNR bundles became more visible when the surrounding CSF and accumulated fluid appeared darker in the selected diffusion-weighted SE-EPI image I_18_ (Fig. 1D), enhancing the contrast between WM in different orientation. On the lesion side, the DREZ and lesion site showed signal intensity changes in diffusion-weighted images, appearing either hypointense (in I_30_) or hyperintense (in I_18_), depending on whether the diffusion gradient direction was parallel or perpendicular to the spinal cord (Fig. 1D). These observation highlights how specific diffusion-weighted contrasts can help differentiate WM fibers in different orientations. In summary, tissue abnormalities in the DNR and DREZ regions were clearly detected and distinguished using a combination of MTC and diffusion-weighted images.

### 3.2 Region-specific changes detected within the spinal cord by multiparametric MRI

Diffusion parametric maps revealed structural changes within the spinal cord, even though the nerve root injury occurred outside the cord. The lesion site in the DNR region (marked by a red cross in Fig. 2) exhibited drastic increased RD and decreased FA. The adjacent

**Figure 2.**
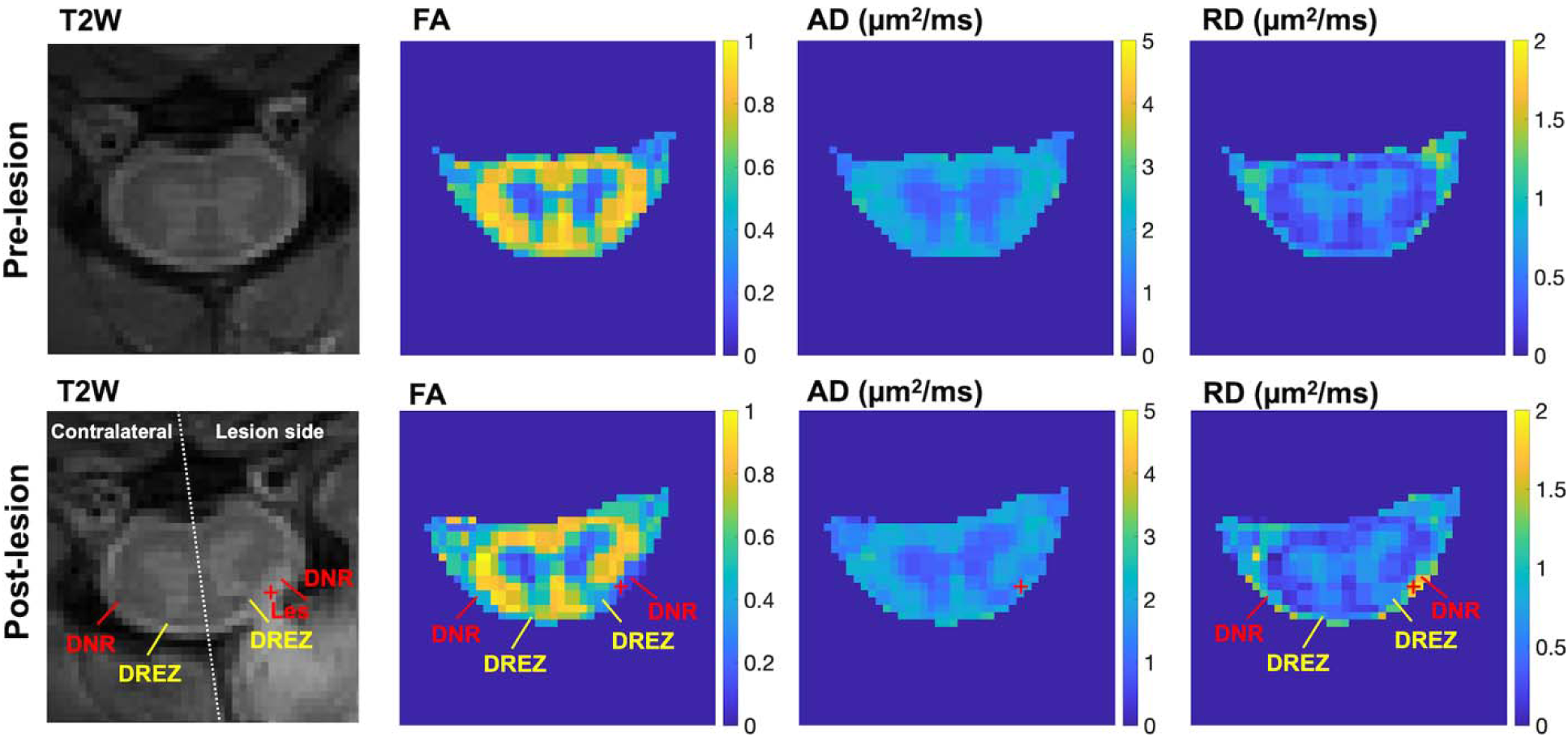
Comparison of diffusion tensor imaging (DTI) derived parametric maps within the selected spinal segment in pre-lesion versus post-lesion conditions. FA (fractional anisotropy), AD (axial diffusivity), RD (radial diffusivity). Red cross indicates the lesion site of dorsal sensory nerve roots. DNR, dorsal nerve roots; DREZ, dorsal root entry zone; Les, lesion site, indicated by symbol ‘+’. Unilateral changes in DNR and DREZ regions are detected and indicated in FA and RD maps acquired post-lesion.

DREZ inside the spinal cord also showed decreased FA and increased RD (yellow arrow in Fig. 2), compared to the contralateral side. Notably, no substantial changes were detected in other ipsilateral regions such as the dorsal horn, dorsal pathway, or lateral pathway. These unilateral changes in diffusion measures were consistently observed at the two spinal segments that underwent nerve root sectioning (slices 2 and 3) when comparing across slices (Fig. 3).

**Figure 3.**
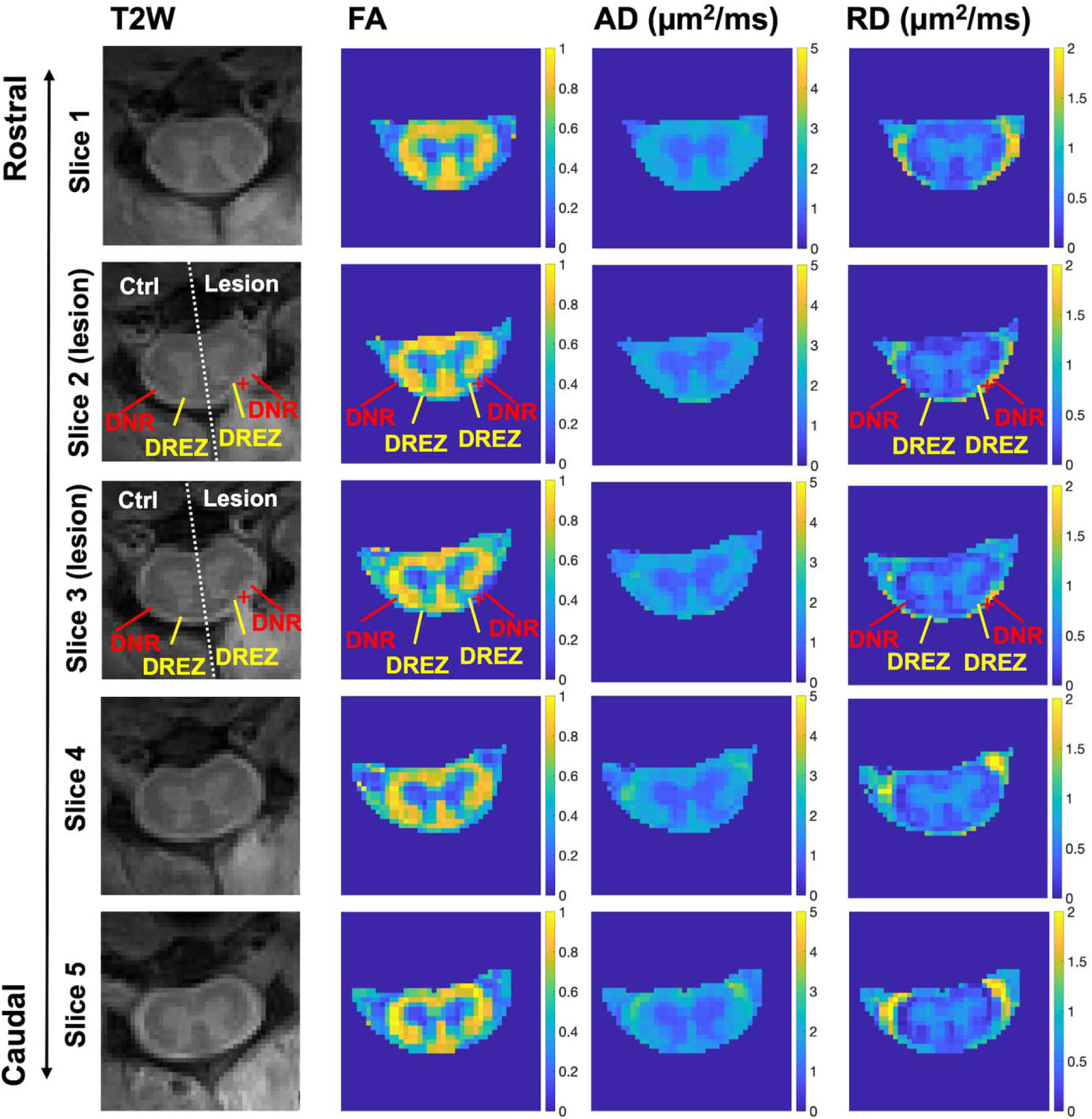
Comparison of DTI-derived parametric maps across spinal cord segments post-lesion. FA, fractional anisotropy; AD, axial diffusivity; RD, radial diffusivity. DNR, dorsal nerve roots; DREZ, dorsal root entry zone. Lesion site is indicated by symbol ‘+’. Ctrl, contralateral side of spinal cord. Unilateral changes in DNR and DREZ regions on the lesion side are detected and indicated in FA and RD maps acquired post-lesion.

PSR maps, derived from qMT, also revealed changes at and near the lesion sites at the two segments underwent sectioning on the lesion side one-week post-injury. At one-week post-injury, the DNR on the lesion side showed reduced PSR values compared to the contralateral side (Fig. 4). A slight PSR decrease was also observed in the ipsilateral DREZ.

**Figure 4.**
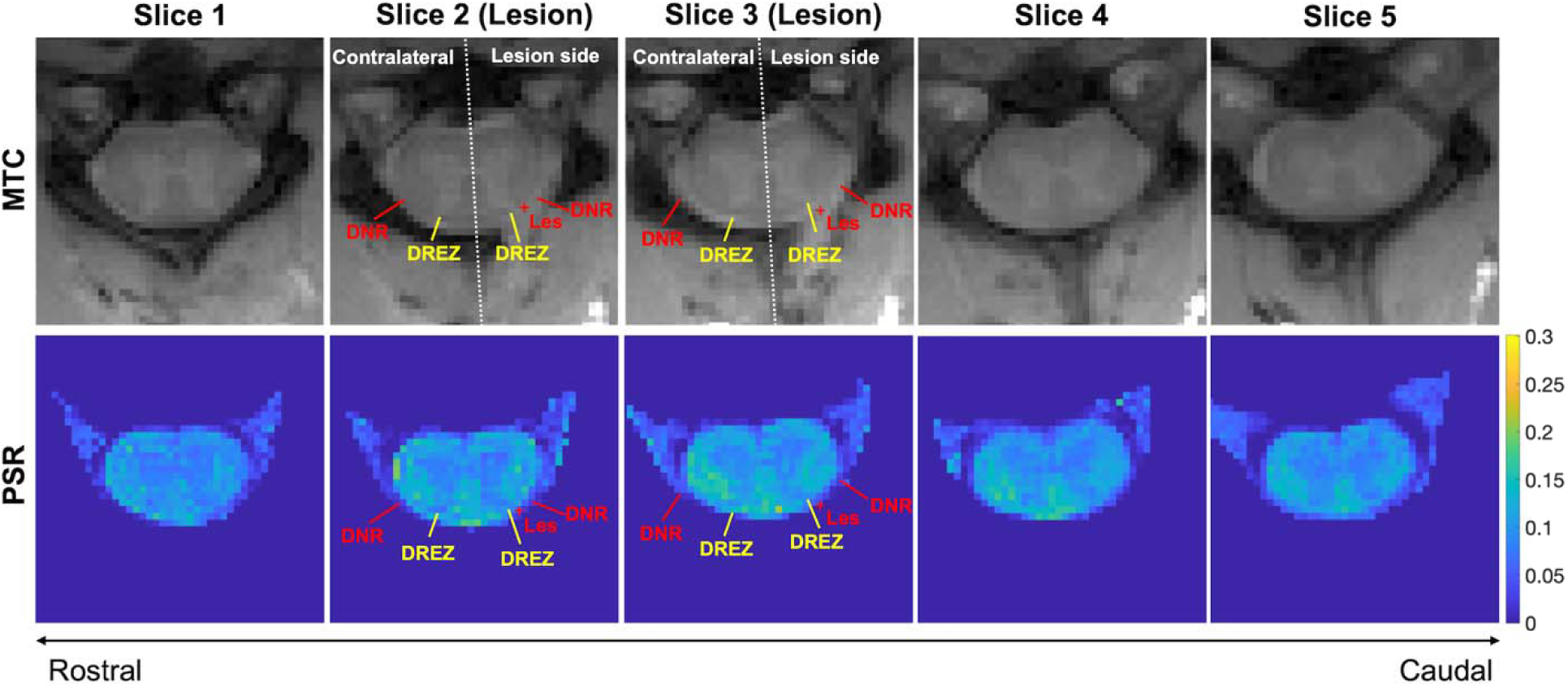
Comparison of pool size ratio (PSR) maps from quantitative magnetization transfer MRI. MTC, magnetization transfer contrast; DNR, dorsal nerve roots; DREZ, dorsal root entry zone; Les, Lesion site, indicated by red symbol ‘+’. Unilateral changes in DNR and DREZ regions are detected and indicated in PSR maps. Results are from a representative subject after dorsal nerve root section.

The Z-spectra of seven ROIs on contralateral and lesion sides were compared (Fig. 5B). Peak amplitudes at different RF offsets were derived from 5-pool fitting of each Z-spectrum to assess CEST and NOE effects (Fig. 5C). The injured DNR (iDNR) and DREZ (iDREZ) showed increased CEST effects around 3.5 ppm RF offset and decreased NOE at -1.6 ppm RF offset (red and blue asterisks, Fig. 5C). The iDNR region also showed a reduced semisolid magnetization transfer effect at 5 ppm RF offset (red double-headed arrow) and diminished NOE effect at -3.5 ppm RF offset (green asterisk), likely due to severe nerve root damage and partial volume effects from fluid accumulation. Other nearby regions, such as the ipsilateral dorsal horn (iDH), showed no evident abnormalities one-week post-injury (Fig. 5B-C).

**Figure 5.**
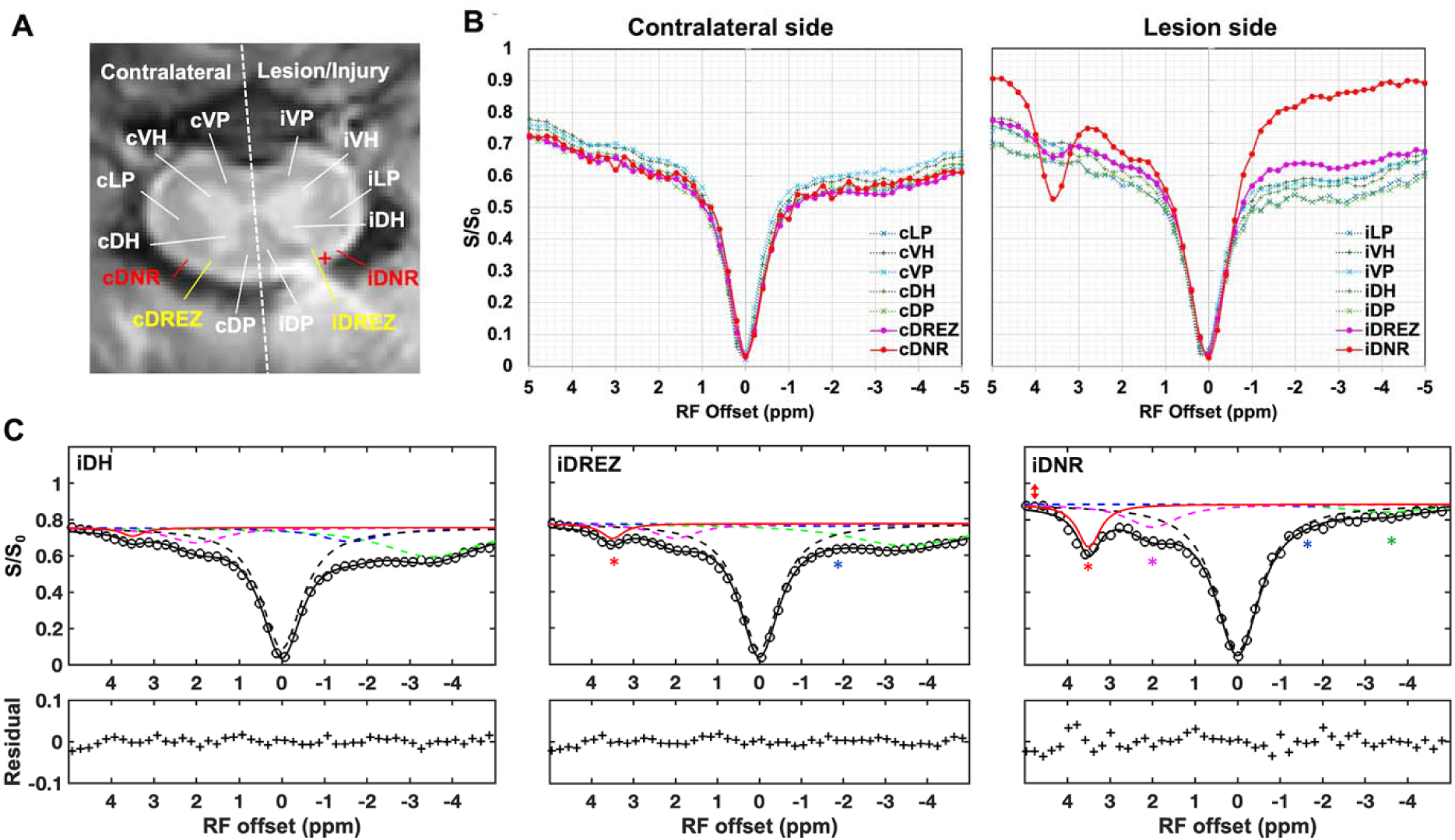
Comparison of averaged regional CEST spectra. (A) Regions of interest. White matter regions include VP (ventral pathway), LP (lateral pathway), and DP (dorsal pathway). Gray matter regions include VH (ventral horn) and DH (dorsal horn). DREZ (dorsal root entry zone); DNR, dorsal nerve roots. The letter “c” and “i” in the ROI labels indicate contralateral and injury side respectively. The red cross indicates the lesion site of DNR. (B) Comparison of the regional spectra on the contralateral and lesion sides in the segments underwent dorsal nerve root injury. (C) The selected regions showing 5-pool Lorentzian fitting results (-3.5, -1.6, 0, 2.0, 3.5 ppm RF offsets). The asterisks indicate significant increase or decrease of amplitudes of the respective fitted peaks. The double arrow indicates semisolid magnetization transfer effect. Results are from a representative subject after lesion.

### 3.3 Group-level comparison of DTI, qMT, and CEST metrics across ROIs

We performed group-level, ROI-based analyses of the MRI measures across seven ROIs (VH, DH, LP, VP, DP, DREZ, and DNR) in pre-lesion and injured spinal cords (Fig. 6). Significant changes in the lesioned DNR were detected in FA (Fig. 6A), PSR (Fig. 6D), CEST at 3.5 ppm (Fig. 6E), and NOE at -1.6 ppm and -3.5 ppm (Fig. 6G-H). Although AD decreased and RD increased in the lesioned DNR (Fig. 6B-C), these changes were not statistically significant (*p* = 0.161 and *p* = 0.065, respectively). Similarly, the increase in CEST at 2.0 ppm was not significant (*p* = 0.232). The DREZ on the lesion side exhibited significant decreases in FA and NOE (-1.6 ppm) and an increase in CEST at 3.5 ppm. None of the other ROIs, including those adjacent to the lesion (e.g. VH, VP, LP, VP and DP), exhibited significant changes at this early post-lesion time point.

**Figure 6.**
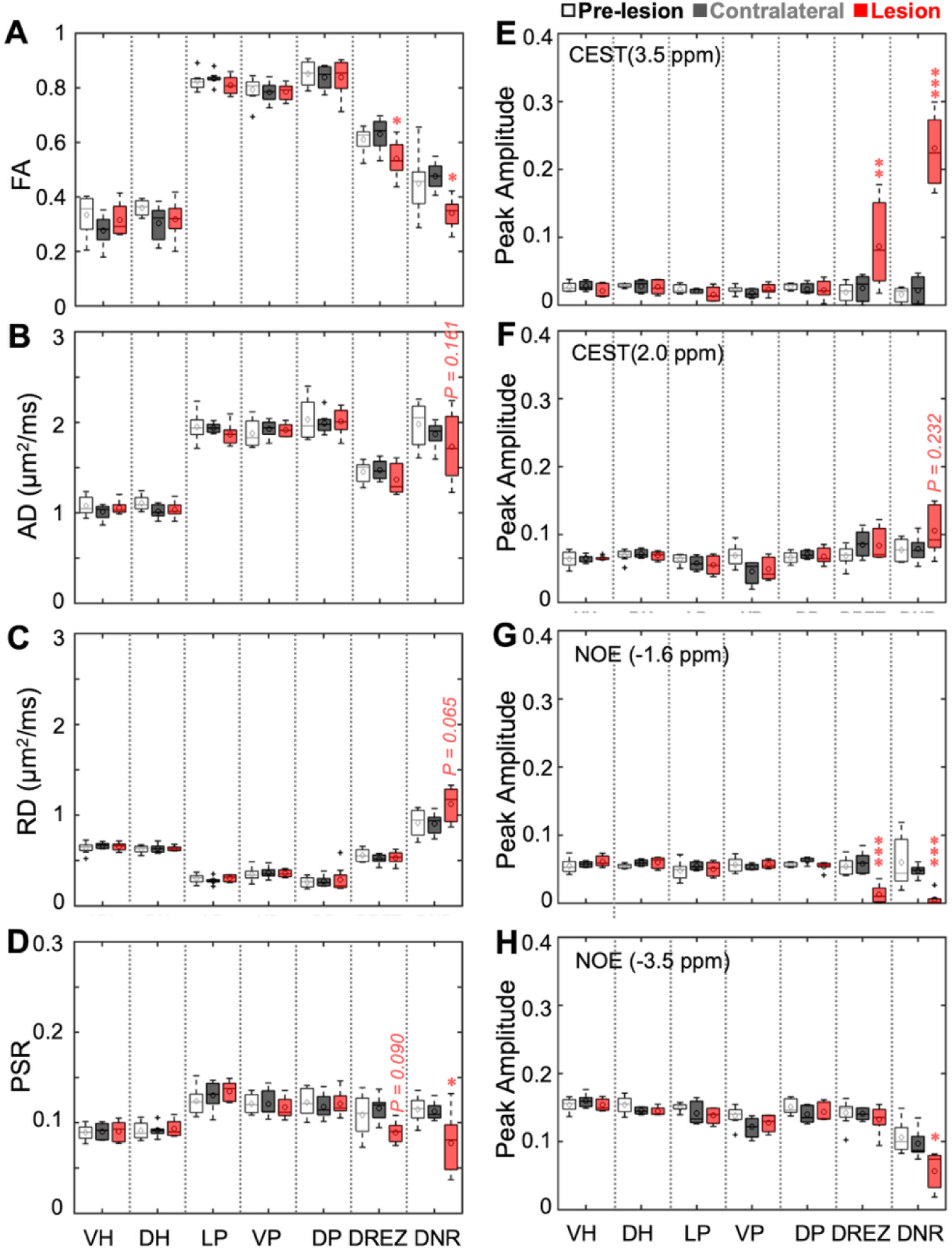
Group level comparison of DTI-, qMT-, and CEST-derived MRI metrics at specific ROIs across all segments (C4 and C5) underwent nerve root injury. Boxplots of MRI measures of non-lesioned contralateral side and lesion side post-lesion with those from respective pre-lesion healthy spinal cord segments (n = 8) in several ROIs: three white matter ROIs (VP: ventral pathway, LP: lateral pathway, DP: dorsal pathway), two gray matter ROIs (VH: ventral horn, DH: dorsal horn), DREZ (dorsal root entry zone), and DNR (dorsal nerve roots). Definitions of these ROIs are illustrated in Figures 1 and 5. (A-C) Diffusion measures FA (fractional anisotropy), AD (axial diffusivity), and RD (radial diffusivity). (D) PSR (pool size ratio) from qMT. (E-H) Peak amplitudes showing the CEST and NOE effects at different RF offsets, including CEST(3.5 ppm), CEST(2.0 ppm), NOE(-1.6 ppm) and NOE(-3. 5 ppm). In the boxplots, middle lines indicate medians and markers indicate mean values. \**p* < 0.05, \*\**p* < 0.01, and \*\*\**p* < 0.001 vs. the corresponding regional values for the normal spinal cords (*Wilcoxon rank sum test*).

### 3.4 Regional correlations between MRI measures

To investigate the relationships among MRI metrics, we performed correlation analyses across ROIs. Figure 7 shows the correlation matrix of all paired comparisons using values from the seven ROIs across the two lesioned segments in four subjects, before and after lesioning (n = 168). Strong correlations (|*r*| > 0.6) were observed between metrics from the same MRI modality, such as positive correlations between FA and AD, and negative correlations between FA and RD. Additionally, CEST at 3.5 ppm was inversely correlated with NOE effects. In addition, PSR from qMT also exhibited strong positive correlations with DTI-derived FA and AD.

**Figure 7.**
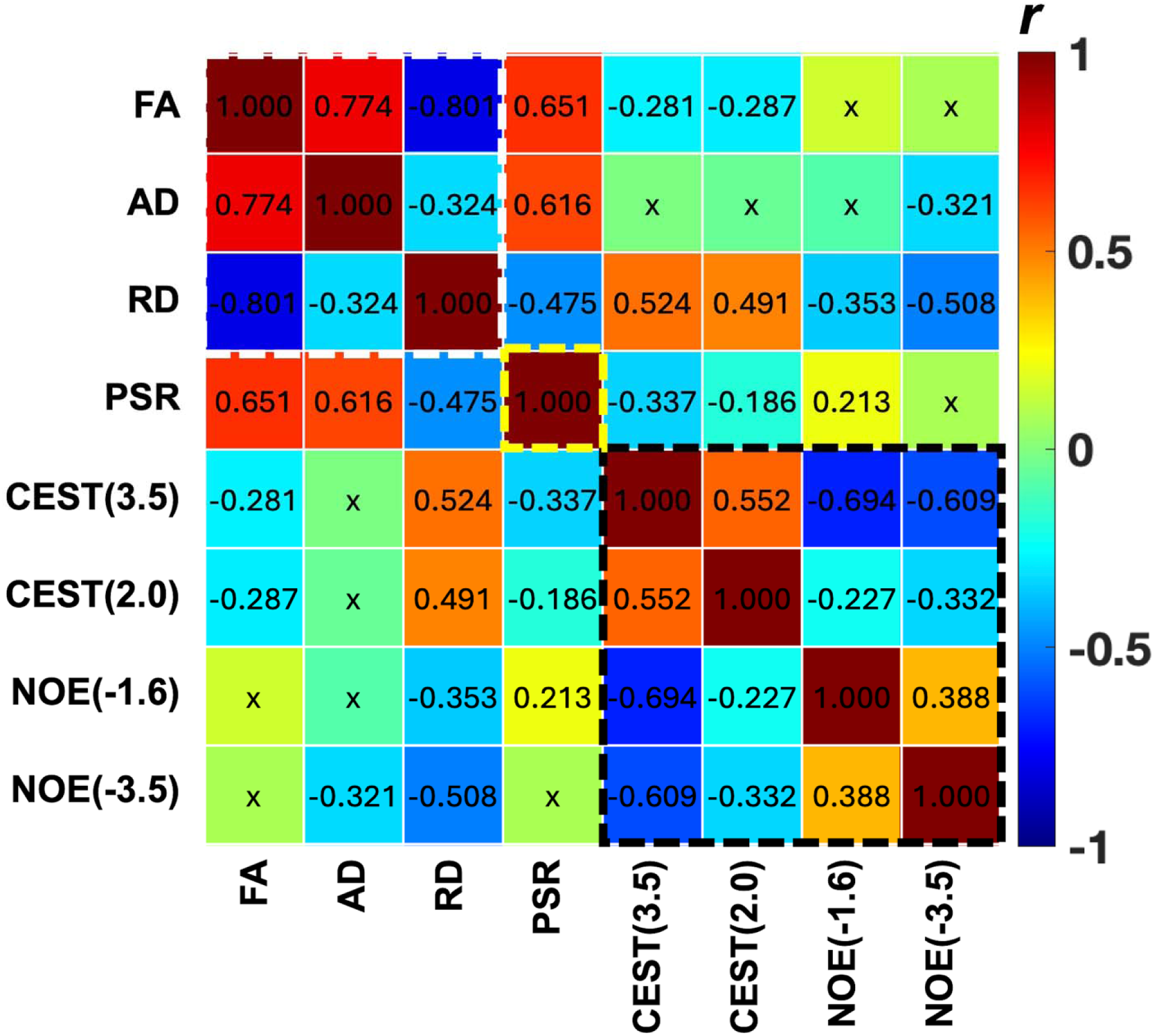
Regional correlations between MRI measures. Correlation coefficients *r* values are shown. ‘x’ indicates correlation that is not significant (*p* > 0.05). Diffusion measures: FA, fractional anisotropy; AD, axial diffusivity; RD, radial diffusivity. PSR, pool size ration from quantitative magnetization transfer. CEST(3.5), CEST(2.0), NOE(-1.6) and NOE(-3.5) are peak amplitudes of peaks at RF offsets 3.5, 2.0, -1.6 and -3.5 ppm from fitting of CEST spectra. Total 168 entries for each metric were included in the correlation analysis.

## 4 DISCUSSION

Multiparametric MRI revealed complex, region-specific pathological changes in the lesioned DNR and adjacent spinal cord tissue after nerve root injury. These changes likely reflect a combination of inflammation, neurotransmitter byproduct release, demyelination, and fiber damage in the regions surrounding the lesion site.

### 4.1 Regional tissue property changes within the spinal cord following dorsal rhizotomy

In the injured DNR region (Fig. 6E-H), increased CEST signals (3.5 ppm and 2.0 ppm), representing amide and amine proton transfer effects, were accompanied by decreased NOE effects at -1.6 ppm and -3.5 ppm, indicative of alterations in both mobile and immobile macromolecular pools. These patterns may arise from the breakdown of neurotransmitters and myelin degradation. The observed increase in CEST at 3.5 ppm in DREZ, along with a decrease in NOE at -1.6 ppm, likely reflects secondary injury, including nerve fiber degeneration and inflammation within the spinal cord caused by damage to the DNR afferents. The elevated amide proton transfer effect may be attributed to a local increase in mobile proteins and peptides where afferent terminals were disrupted, or to acidification of the tissue microenvironment. The NOE at -1.6 ppm, associated with aliphatic protons in lipids and membrane proteins, diminishes in response to demyelination and tissue damage. Collectively, changes in CEST and NOE suggest a disrupted local chemical environment associated with inflammation, apoptosis, and secondary axonal degeneration in the DNR and DREZ regions. Decreased FA in these regions further supports the presence of fiber degeneration and demyelination. Corresponding reductions in AD and increases in RD in the DNR align with these changes. PSR from qMT, which is sensitive to myelin content, also decreased in DNR and DREZ, reinforcing evidence for demyelination.

Significant increases in CEST (3.5 ppm) and decreases in NOE (-1.6 ppm) in the DREZ indicate early inflammatory responses and secondary injury. These biochemical changes coincide with decreases in PSR and FA, further supporting the occurrence of demyelination and fiber disruption. We hypothesize that focal unilateral dorsal nerve root sectioning would initiate afferent fiber degeneration, which could lead to secondary injury, causing fiber damage that extends along the central branch of the injured nerve roots bundles and into entering zones.

### 4.2 Advantage of multiparametric MRI in detecting early-stage changes following nerve root sectioning

Our previous studies have shown that multiparametric MRI, incorporating DTI, qMT and CEST, is sensitive to pathological changes in both GM and WM following traumatic SCI.^11, 12, 14–18^ These techniques enable comprehensive assessments of injury progression and recovery. In the present study, we provide MRI evidence that transecting of dorsal nerve roots outside the spinal cord leads to measurable changes in WM structural integrity and tissue neurochemistry within one week post injury. This finding holds clinical significance, as selective dorsal rhizotomy is a neurosurgical procedure to alleviate pathological sensory inputs in conditions such as cerebral palsy, neuropathic pain, and SCI-related spasticity.^2^ Our results suggest that multiparametric MRI provides biomarkers that can serve as noninvasive indicators of spinal cord pathology associated with dorsal nerve root injury.

Specifically, dorsal root injury caused detectable alterations in both central branches of the dorsal nerves and the adjacent DREZ region. Among all MRI metrics, CEST (3.5 ppm) and NOE (-1.6 ppm) showed the most significant changes (smallest *p*-values) in the DNR and DREZ regions one-week post-lesion (Fig. 6). FA was also reduced in both regions, while PSR showed a pronounced decrease in the DNR, consistent with demyelination of the central afferent terminals. A modest PSR decrease in the DREZ suggests mild demyelination at the nerve entry zone. Together, reductions in PSR and FA in the lesioned DNR and adjacent region DREZ support the presence of secondary axonal degeneration and demyelination extending from the peripheral injury into the intact spinal cord. The accompanying CEST and NOE changes confirm inflammation and altered biochemical composition. These observations highlight the utility of multiparametric MRI in detecting subtle, localized spinal cord pathology soon after peripheral nerve injury.

The ability to noninvasively monitor such delicate regional alterations is critical for both preclinical and clinical studies, especially when investigating secondary effects arising from upstream dorsal sensory disruptions. Detecting these pathological changes at the central terminals of sparsely distributed nerve roots is particularly challenging, as they may span one to two spinal segments rostrally and caudally. In this context, multiparametric MRI offers a powerful tool to monitor structural and neurochemical changes and interpret their functional implication following nerve root injury.

### 4.3 Interpretation of inter-pool correlations

Strong correlations were observed between metrics from the same imaging modality, suggesting that these parameters covary and provide complementary insights into different aspects of spinal cord pathology. The high correlation between amide and amine CEST pools implies a shared origin in mobile proteins and peptides under normal conditions. After injury, the degradation of macromolecules and release of small peptides and amino acids likely contributed to the observed negative correlations between NOE and CEST metrics.

Among DTI and qMT metrics, RD showed the strongest correlation with CEST (3.5 ppm), indicating a close association with DNR degeneration and demyelination. Inflammation may influence RD more than AD, especially given the orientation of nerve roots. While partial volume effects from surrounding CSF can attenuate RD changes, the observed trends remain informative. Post-lesion decreases in PSR and FA were consistent with WM fiber damage, and these two measures were strongly positively correlated. This relationship reinforces their joint utility as markers of subtle WM abnormalities within the intact spinal cord. PSR also correlated slightly more strongly with CEST (3.5 ppm) than FA (Fig. 7), possibly because PSR directly reflects the fraction of intact macromolecular pools, while CEST (3.5 ppm) highlights biochemical changes from fiber degeneration. FA, while significantly correlated with PSR due to the organized fiber structure of the spinal cord, does not directly measure fiber fraction and may be less relevant in abnormal tissues with complex or disrupted fiber organization. Despite these distinctions, both PSR and FA reliably decreased in regions of fiber loss and exhibited strong correlation with each other, supporting their potential as imaging biomarkers for nerve injury and degeneration.

## 5 CONCLUSIONS

This study demonstrates that multiparametric MRI can detect localized structural and biochemical changes within the intact spinal cord following unilateral dorsal root nerve section in a non-human primate model of dorsal rhizotomy. One-week post-injury, MRI revealed early signs of WM fiber degeneration, demyelination, and neuroinflammation at the central terminals of the dorsal root and the adjacent DREZ. These findings suggest that peripheral afferent injury can induce secondary spinal cord pathology. Multiparametric MRI offers a sensitive, noninvasive approach for characterizing these changes and has potential for monitoring injury progression and evaluating therapeutic interventions.

## ACKNOWLEDGEMENTS

We thank Mrs. Chaohui Tang and Mr. Fuxue Xin of the Vanderbilt University Institute of Imaging Science for their assistance in animal preparation and care during MRI data collection. We also thank Drs. Ming Lu and Xinqiang Yan for customizing coils for cervical spinal cord imaging. This study is supported by DOD grant W81XWH-17-1-0304, and NIH grant NS092961.

## Notes

### Competing Interest Statement

The authors have declared no competing interest.

